# Sensitivities in associating land-system archetypes with sustainability metrics: Insights from simulations

**DOI:** 10.1101/2024.08.01.606212

**Authors:** Varun Varma, Paul M Evans, Luca Bütikofer, Cecily E D Goodwin, Richard F Pywell, John W Redhead, Jonathan Storkey, James M Bullock, Andrew Mead

## Abstract

Archetypes of land- and socio-ecological systems, generated using unsupervised classification methods, enable the assimilation of complex environmental and socio-economic information. Such simplification has considerable potential to feed into decision support systems for sustainability planning. But, the usefulness of archetypes depends on how well they relate to sustainability criteria, such as ecosystem service (ES) delivery, that are external to the input datasets employed for archetype generation. Sensitivities in such post-hoc association analyses, and the associated utility of the archetype framework in a decision support context, remain unexplored. Here we emulated post-hoc association analysis procedures using simulated socio-ecological datasets and ES response variables. Our simulations revealed a substantial influence on analysis performance from (1) the number of variables used as inputs in archetype generation, (2) the correlation structure of input datasets, (3) the type and distribution of input variables, and (4) the functional form (linear or non-linear) characterising the relationship between ES variables and their predictors. We observed near-identical performance when archetypes were generated using K-means clustering and Self-Organising Maps (SOMs) – two commonly used archetype classification methods. Further, better archetype classifier performance did not guarantee better discrimination of ES value distributions between archetypes. Our results suggest that designing a framework to generate archetypes for sustainability planning, and the selected methodological choices therein, should place greater emphasis on what the archetypes will be used for in downstream analyses, and not focus solely on archetype classifier performance. This would better ensure the identification of archetypes adaptable to a diverse array of sustainability indicators and sufficiently robust for monitoring decision outcomes over time.

## 1. Introduction

Land-system typologies are increasingly being proposed as a way to organise highly dimensional datasets, including varied data types in the context of supporting decision making towards sustainability goals (Václavík *et al*., 2013; Oberlack *et al*., 2019). These typologies, or land-system archetypes, capture coupled human-nature interactions, or socio-ecological systems, by identifying recurring patterns in variables describing landscapes and/or processes that operate within them (Ostrom, 2009; Eisenack, 2012; Oberlack *et al*., 2016). Dividing a landscape into archetypes usually involves unsupervised classification, where classes or archetypes represent parcels of land that share similarities in input variable values. The input variables can encode information on fixed (e.g., elevation) or easily-modifiable (e.g., land management) properties, and hence, can represent existing associations between these variables and indicators of sustainability goals, as well as provide information on potential pathways to enhance indicators of interest (van Asselen and Verburg, 2012; Cullum *et al*., 2017; Tieskens *et al*., 2017; Goodwin *et al*., 2022). This approach explicitly assumes that there may be multiple, context-dependent models of associations between landscape properties and indicators of sustainability (Young *et al*., 2006; Eisenack *et al*., 2019; Oberlack *et al*., 2019). Hence, it aims to operate at an intermediate level of abstraction, where the pitfalls of gross generalisations over a landscape and the highly individual nature of case studies can be overcome (Bureau *et al*., 2012; Frey and Cox, 2015; Cullum *et al*., 2017; Frey, 2017; Oberlack *et al*., 2019; Pratzer *et al*., 2024). The simplification of complex data associations and interactions to archetypes can serve as a valuable communication tool for stakeholders, while also providing a context-dependent framework for decision making. However, a degree of information loss is inevitable when observations are categorised in this way (Cox, 1957; Steel *et al*., 2013; Busch, 2021). This may be an acceptable trade-off, given the potential of such analyses. Nevertheless, we need to assess how methodological choices affect information loss, and how this could be minimised.

Analyses that develop and/or use land-system and socio-ecological archetypes can broadly be divided into three types - descriptive, internal associations, post-hoc associations - depending on how they incorporate potential predictor and response variables of a system. Descriptive analyses are where variables on land-system properties, or potential predictors, are clustered to form archetypes. Potential response variables are not included (or variables are not explicitly identified as responses) in the clustering process, though the resulting archetypes form a framework within which response variables could be sampled from and analysed at a later stage (Beckmann *et al*., 2022). For example, Václavík *et al*. (2013) utilised Self-Organising Maps (SOMs) – a popular unsupervised classification algorithm in archetype analyses (Lek and Guégan, 2000; Chon, 2011; Levers *et al*., 2018; Sietz *et al*., 2019) – to develop a schema of 12 global land-system archetypes. These archetypes describe recurring patterns across 32 variables capturing properties of land-use, environmental conditions and socio-ecological factors. SOMs have also been used in more regional analyses, such as the development of agri-environmental archetypes for Europe (Beckmann *et al*., 2022), and non-nested tiered archetypes of agricultural systems in Great Britain (Goodwin *et al*., 2022). The latter provides a multi-layered contextual classification of landscape pixels into archetypes. Conceptually similar analyses have also been applied to global datasets identifying anthropogenic biomes (Ellis and Ramankutty, 2008; Ellis *et al*., 2010), as well as to track changes in the spatial configuration of archetypes, and land-use intensity within archetypes over time (Ellis *et al*., 2010; Levers *et al*., 2018; Mengxue *et al*., 2022).

Internal association analyses are where potential predictor variables, and one or more response variables of interest are classified into archetypes that represent recurring patterns in the distribution of predictors and responses. The co-occurring patterns in the distributions are then used to infer associations between responses and archetypes, and implicitly to the predictor variables that characterise each archetype. This thematic classification allows for inferences on where and which management strategies co-occur positively with improving sustainability indicators, identify regions that are vulnerable, and hence, where transfer of strategies would result in desirable outcomes (e.g., Václavík *et al*., 2016; Sietz *et al*., 2017; Rocha *et al*., 2020; Obringer and White, 2023).

The third analysis type we term post-hoc association analysis, where archetypes of land-systems are formulated as in descriptive analyses (type 1), and response variables of interest are then associated with the archetypes. There can be varied approaches to linking responses to archetypes, including simple spatial intersections (Adenle and Ifejika Speranza, 2021), multiple regression approaches (Nair *et al*., 2016) and by combining classification of multiple predictors and responses (Oh *et al*., 2007; Tison *et al*., 2007). Post-hoc association analyses constitute an extension of descriptive analyses (type 1) and differ from internal association analyses (type 2) in that response variables remain external to data processing that defines the archetypes. This feature provides an important advantage in a decision support system, as responses of interest could change over time, either due to availability of improved datasets (Nguyen *et al*., 2022), new or improved ways of measuring responses of interest (Johnson *et al*., 2021; Lynggaard *et al*., 2024), or identification of new priorities and trade-offs as sustainability science and our understanding of ecosystems improve. Incorporating modified or new response variables within post-hoc association analyses would not require the redefinition of archetypes that is likely required with internal associations analyses.

While descriptive and internal association analysis methods can benefit from guidelines to objectively assess validity (Piemontese *et al*., 2022), methodological sensitivities that could impact on the usefulness of post-hoc associations analyses remain unexplored. The characteristics of input datasets, such as the number and type of variables, along with the nature of inter-variable relationships, are well-established determinants of statistical model and classifier behaviour (Kiang, 2003; Li and Lin, 2014; Petitpierre *et al*., 2017), and could influence the analytical performance of archetype analyses. Here, we use simulations to investigate the sensitivities of archetype analysis in detecting post-hoc associations between sustainability indicators and archetypes. Specifically, we generated datasets of predictors and responses (emulating ecosystem services; ESs) to explore how commonly used methods to derive archetypes (K-means clustering and SOMs), the structure of the input data (number of variables, correlation structure and variable types) and the nature of the relationship between ESs and their driver variables (linear and non-linear) influence the potential to detect differences in sustainability indicators between archetypes.

## 2. Methods

The broad logic of the simulations was as follows: we simulated input datasets of 10,000 observations (rows) comprising multiple variables or columns. These represented data structures that researchers might ordinarily assemble to study a landscape, i.e., by compiling environmental, land-use, land management and socio-economic variables. Though our simulations were not spatially explicit, one could consider each observation in the input dataset as a pixel in the landscape. We then assigned each observation an ES value by simulating a deterministic relationship (perfectly predicted, and without error) with a single variable within the input dataset. While a real-world ES likely involves multiple predictors, this simplification allowed us to embed a strong, noise-free signal within the data. The extent to which subsequent analysis recovered this embedded signal could then be used as a metric to assess its analytical performance. As we find, there are numerous challenges when detecting this simple signal. Hence, introducing further complexity within simulations (e.g., multiple predictors for a single ES, multiple ESs, etc.) would not be useful for the purpose of this study.

The input dataset (excluding the simulated ES) was then subjected to unsupervised classification to generate a pre-determined number of archetypes using SOMs or K-means clustering. This resulted in each observation being assigned an archetype identity. For each observation in the input dataset, the Euclidean distance to its assigned archetype centroid (cluster centroid) was computed. The median distance of observations to their respective archetype centroid provides a simple measure of how well the classifier performed in identifying distinct archetypes (Wehrens and Buydens, 2007; Václavík *et al*., 2013), where larger values indicate poorer performance. To determine the extent to which differences in simulated ES values could be detected between archetypes, ES values were sub-setted by archetype identity, and descriptors of ES distributions within archetypes were computed. Specifically, we estimated the first (Q1) and third (Q3) quartiles of within-archetype ES value distributions. Pairwise comparisons of ES inter-quartile ranges between archetypes were performed to estimate a ‘detectability’ score (as a proportion). For example, if observations in the input dataset were classified into eight archetypes, then ES values (one value for every observation) could be sub-setted into eight distributions. There are 28 possible ways of comparing pairs of the eight distributions, and detectability represented the proportion of these 28 pairwise comparisons where ES inter-quartile ranges did not overlap.

We modified properties of the input dataset, the nature of ES-input variable relationships and archetype generation parameters to explore the sensitivities in identifying post-hoc associations between archetypes and ESs. As the number of possible modifications and their combinations were exceedingly large, we partially stratified simulations into four sets (Table 1). All simulations and analyses were run using R version 4.2 (R Core Team, 2023) and a detailed description of the simulation design for each set is as follows:

**Table 1.**
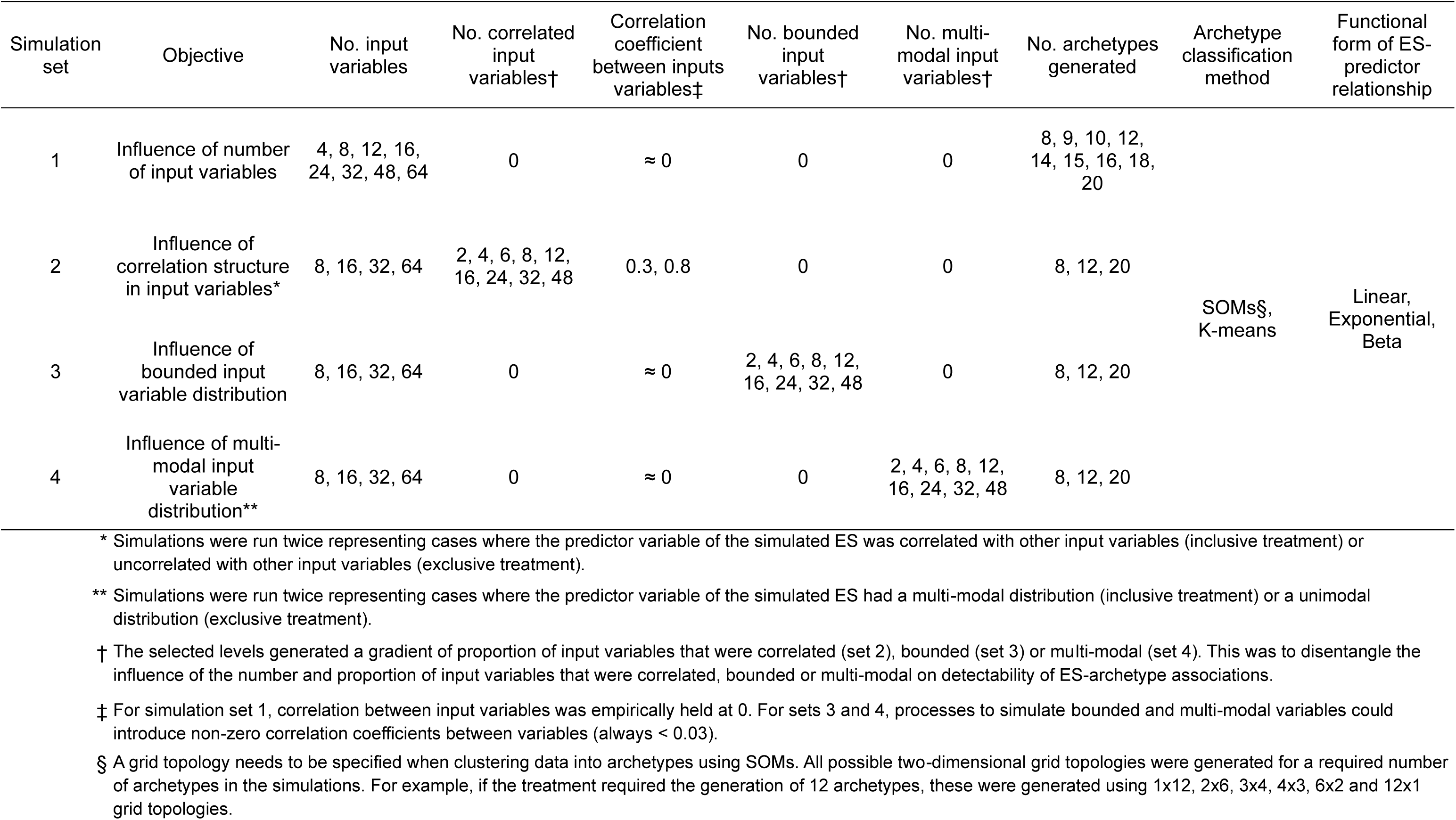
Summary of simulation sets identifying objectives of each set and ‘treatment’ applied. Detailed description of the methodology for generating each simulation set, including the choice of parameter values, is provided in the Methods sections 2.1 to 2.4.

### 2.1. Set 1 – effects of number of input variables, number of archetypes and classification method

Set 1 simulations investigate the influence of increasing the number of variables in an input dataset, the number of archetypes the dataset is classified into and classification methodology on the detectability of post-hoc associations between archetypes and ESs. We simulated input datasets with differing numbers of variables (eight levels of dataset size: 4, 8, 12, 16, 24, 32, 48 and 64 variables). The datasets were generated using the R package *faux* (version 1.2.1; DeBruine, 2023), such that each variable was normally distributed (mean = 0; standard deviation = 1), and none of the variables were correlated to each other (correlation coefficients were empirically forced to zero). One variable in each dataset was used to define a simulated ES using a linear, exponential or beta function (see below – ES functions). We classified the observations of the input dataset into different numbers of archetypes (nine levels for number of archetypes: 8, 9, 10, 12, 14, 15, 16, 18 and 20 archetypes) using SOMs (R package *kohonen* (version 3.0.12; Wehrens and Kruisselbrink, 2018)) and K-means clustering. Constructing SOMs for a given number of archetypes requires users to specify a grid topology. We used all possible rectangular representations of grid topologies for our candidate number of archetypes (e.g., for eight archetypes we used all four possible grid topologies 1x8, 2x4, 4x2, 8x1; for 12 archetypes we used all six possible grid topologies 1x12, 2x6, 3x4, 4x3, 6x2, 12x1). For the nine treatment levels of number of archetypes, this corresponded to 42 ‘number of archetype x grid topology’ configurations for SOMs, and nine K-means clustering scenarios. Fifty replicate datasets were generated for each dataset size (i.e., 400 datasets across eight levels of dataset size). Each dataset was used to simulate ES values using the three functional forms. Datasets were then processed to generate differing number of archetypes (nine levels) using the 51 clustering scenarios (42 SOM scenarios + nine K-means scenarios).

### 2.2. Set 2 – effects of correlated input variables

Here, simulations investigated the influence of correlation structure within input datasets on the detectability of post-hoc associations between archetypes and ESs. Datasets of four different sizes were generated (dataset size levels: 8, 16, 32 and 64 variables). In contrast to datasets generated in set 1 simulations, a subset of variables within datasets here were correlated at two different levels (correlation levels: Pearson’s correlation coefficients of 0.3 and 0.8). Simulated datasets with eight variables included 2, 4 or 6 correlated variables; datasets with 16 variables included 2, 4, 6, 8 or 12 correlated variables; datasets with 32 variables included 2, 4, 6, 8, 12, 16 or 24 correlated variables and datasets with 64 variables included 2, 4, 6, 8, 12, 16, 24, 32 or 48 correlated variables. In each dataset, correlated variables shared the same correlation coefficient, e.g., in a simulated dataset of 16 variables with four correlated variables, all four shared the same correlation coefficient with each other (0.3 or 0.8), and correlation coefficients of uncorrelated variables in the dataset were empirically forced to zero. This design provides treatment combinations to disentangle the influence of the number and proportion of correlated variables on detectability. Further, corresponding dataset size levels from set 1 simulations, where all variables within input datasets were uncorrelated, serve as a control to assess the overall effect of introducing correlation structure into input datasets.

As with set 1 simulations, ES values were generated (using linear, exponential or beta functions) as a function of one input variable within the dataset. However, we introduced two treatments here, where the ES predictor variable was either a member or a non-member of the correlated subset of variables within the input dataset (i.e., two membership levels: inclusive and exclusive). To economise on computation time, we reduced the number of archetypes that input datasets were classified into to three levels (8, 12 and 20 archetypes). Again, archetypes were generated using SOMs (using all possible rectangular grid configurations given the number of required archetypes) and K-means clustering. We generated 50 datasets of each combination of [(dataset size-number of correlated variables) x correlation level x membership level] {i.e., 50 x [(24) x 2 x 2] = 4800 datasets}. Datasets were then processed to generate differing number of archetypes (three levels) using the 19 clustering scenarios (16 SOM scenarios + three K-means scenarios).

### 2.3. Set 3 – effects of bounded input variables

Bounded variables are frequently encountered in ecological datasets, e.g., when classes of land-use and land-cover datasets at finer resolutions than the target analysis resolution are coarsened to percent or proportion cover within pixels, or when composition of soil is expressed as percentages. To assess the influence of such variables on the detectability of archetype-ES associations, we generated datasets – using the same methodology as used in set 1 simulations – of four different sizes (dataset size levels: 8, 16, 32 and 64 variables), where all variables were uncorrelated. We then truncated the distributions of a subset of variables in each dataset to generate bounded distributions with symmetrical peaks at the ends of the distribution. We varied the number of variables within datasets with such distributions, such that datasets with eight variables included 2, 4 or 6 bounded variables; datasets with 16 variables included 2, 4, 6, 8 or 12 bounded variables; datasets with 32 variables included 2, 4, 6, 8, 12, 16 or 24 bounded variables and datasets with 64 variables included 2, 4, 6, 8, 12, 16, 24, 32 or 48 bounded variables. As with set 2 simulations, this design allowed us to isolate the influence of the number and proportion of bounded variables on detectability, and corresponding dataset size levels from set 1 simulations acted as controls to assess the overall effect of incorporating bounded variables into input datasets. When generating ES values for each dataset, we restricted the deterministic predictor variables to only those which did not have bounded distributions. Similar to set 2 simulations, input datasets were only classified at three levels (8, 12 and 20 archetypes) using SOMs (and all possible rectangular grid configurations for a given number of required archetypes) and K-means clustering. We generated 50 replicate datasets for each ‘dataset size-number of bounded variables’ combination (i.e., 50 x 24 = 1200 datasets). Datasets were then processed to generate differing number of archetypes (three levels) using the 19 clustering scenarios (16 SOM scenarios + three K-means scenarios).

### 2.4. Set 4 – effects of multi-modal input variables

Variables with multi-modal distributions within a study landscape could arise through various mechanisms, such as when study landscapes contain geographic discontinuities or barriers. To assess the sensitivity of archetype-ES associations to such variables, we generated datasets of four different sizes (dataset size levels: 8, 16, 32 and 64 variables) using the same methodology as in set 1 simulations. We then replaced subsets of variables with trimodal distributions containing symmetrical peaks. Each of these variables were simulated by appending random values drawn from normal distributions with means of -2, 0 and 2, and standard deviations of 0.5 resulting in data ranges similar to the remaining unaltered variables within the simulated dataset. The values of each trimodal variable were randomised to break any correlation structure that may have been inadvertently generated. Datasets with eight variables included 2, 4 or 6 trimodal variables; datasets with 16 variables included 2, 4, 6, 8 or 12 trimodal variables; datasets with 32 variables included 2, 4, 6, 8, 12, 16 or 24 trimodal variables and datasets with 64 variables included 2, 4, 6, 8, 12, 16, 24, 32 or 48 trimodal variables. We restricted our simulations to only include trimodal variables for simplicity as there are countless variations that could have been simulated. As with set 2 simulations, we generated ES values (using linear, exponential and beta functions) along two ‘membership’ treatments, where the ES predictor was either a unimodal (exclusive treatment) or trimodal variable (inclusive treatment) within the input dataset. Input datasets were then classified into to three levels of number of archetypes (8, 12 and 20 archetypes) using SOMs (and all possible rectangular grid configurations for a given number of required archetypes) and K-means clustering. We generated 50 replicate datasets for each combination of [(dataset size-number of trimodal variables) x membership level] {i.e., 50 x [(24) x 2] = 2400 datasets}. Datasets were then processed to generate differing number of archetypes (three levels) using the 19 clustering scenarios (16 SOM scenarios + three K-means scenarios).

#### ES functions and parameter values used

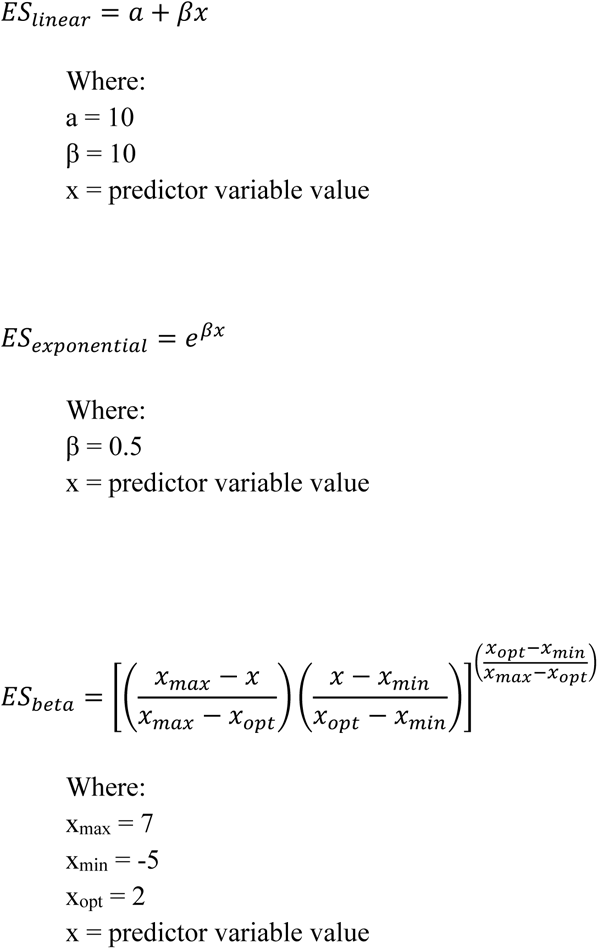

## 3. Results

Our design generated 10,800 unique treatment combinations. Due to its size, we identify key findings below, and present a complete set of summary metrics for each treatment combination in supplementary file SF1.

### 3.1. Set 1 – Influence of number of input variables

Simulations from set 1 (where all input variables were uncorrelated; see Table 1) showed that detectability of differences in a simulated ES between archetypes declined with an increase in the number of variables within the input dataset used to derive the archetypes (Fig 1a, b). Detectability was similar regardless of archetypes being derived using SOMs (and all grid topologies for a given number of archetypes) or K-means clustering. Clustering performance also showed no difference between the two methods, as the increase in median distance to archetype centroids with number of input variables was similar for both methods (Supplementary Fig S1). Loss of detectability was partially mitigated if a greater number of archetypes were generated. For example, detectability was greater when we generated 20 (Fig 1b) compared to eight archetypes (Fig 1a). Also, detectability tended towards zero with a greater number of input variables (> 32 variables) for 20 archetypes compared to > 16 variables for 8 archetypes. Unsurprisingly, the functional form of the relationship between the simulated ES and input variable modified detectability. The linear relationship showed greater detectability, and while the exponential relationship (representing a weak non-linearity) showed no difference in detectability (not shown), the beta relationship (Fig 1c and 1d) showed reduced detectability compared to the linear relationship (Fig 1a and 1b).

**Fig 1.**
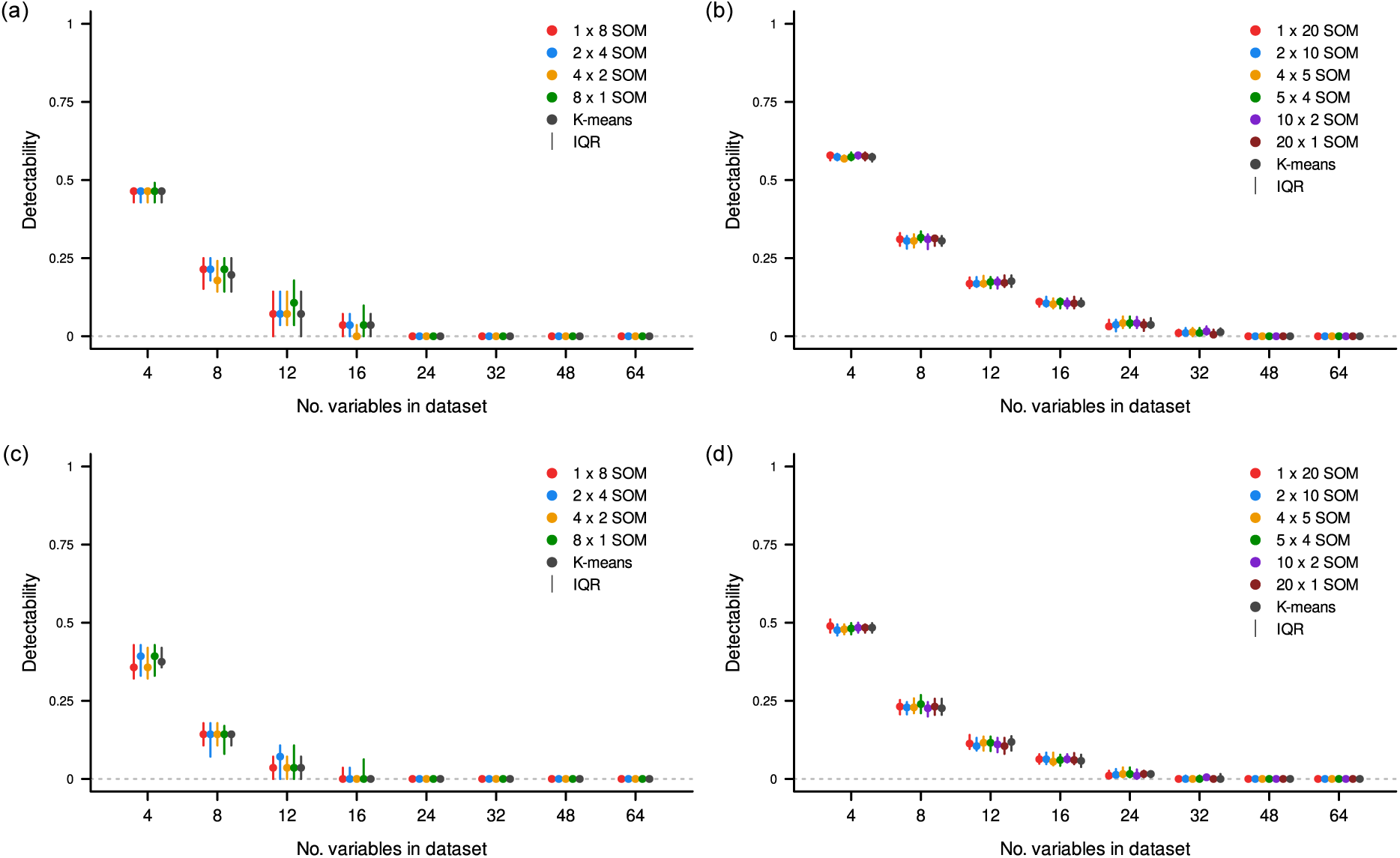
Change in detectability of differences in ecosystem service (ES) values between archetypes generated using an increasing number of input variables. These results are for simulation set 1 (see Table 1) where input variables were not correlated with each other. (a, c) Eight archetypes were generated using all four possible grid topology configurations of Self-Organising Maps (SOMs) or K-means clustering. (b, d) Twenty archetypes were generated using all six possible grid topology configurations of SOMs or K-means clustering. A linear relationship with one variable in the input dataset was used to simulate ES values in (a) and (b), and a beta function was used in (c) and (d). Points represent median detectability from 50 replicates, and vertical lines represent detectability inter-quartile ranges (IQR).

### 3.2. Set 2 – Influence of correlation structure in input variables

Correlated variables within datasets (simulation set 2) had a substantial, yet nuanced, influence on detectability depending on the strength of the correlations, the number and proportion of correlated variables, and if the ES driver variable was amongst the subset of correlated variables. As with set 1 simulations, detectability improved when a greater number of archetypes were generated, declined with a non-linear relationship between ES and input variables, and was not influenced by methodology used (SOM or K-means) to develop archetypes. For clarity, we used the case of 20 archetypes generated using K-means clustering to illustrate key findings in Fig 2. When there were weak correlations between input variables (Fig 2a) detectability improved in datasets where over half the variables were correlated with each other, but then diminished as dataset size (number of variables) increased. When the dataset contained weak correlations, and when the ES driver variable was amongst the correlated variables (inclusive correlations), detectability also reduced with an increase in dataset size, but consistently remained higher compared to datasets with no correlations. Where correlations between input variables were strong (Fig 2b), detectability in exclusive datasets (i.e., where the ES driver variable was not correlated with other input variables) was further improved when half or more of the variables were correlated, and again, reduced with an increase in dataset size. With inclusive correlations, detectability was substantially higher compared to uncorrelated datasets and datasets with weak correlations. The effect of input dataset correlation structure on detectability was strongly influenced by the interaction between the number of correlated variables and the proportion of variables they comprised within the input dataset. When considering inclusive correlations, detectability increased as fewer variables accounted for a greater proportion of correlated variables within datasets (Supplementary Fig S2).

**Fig 2.**
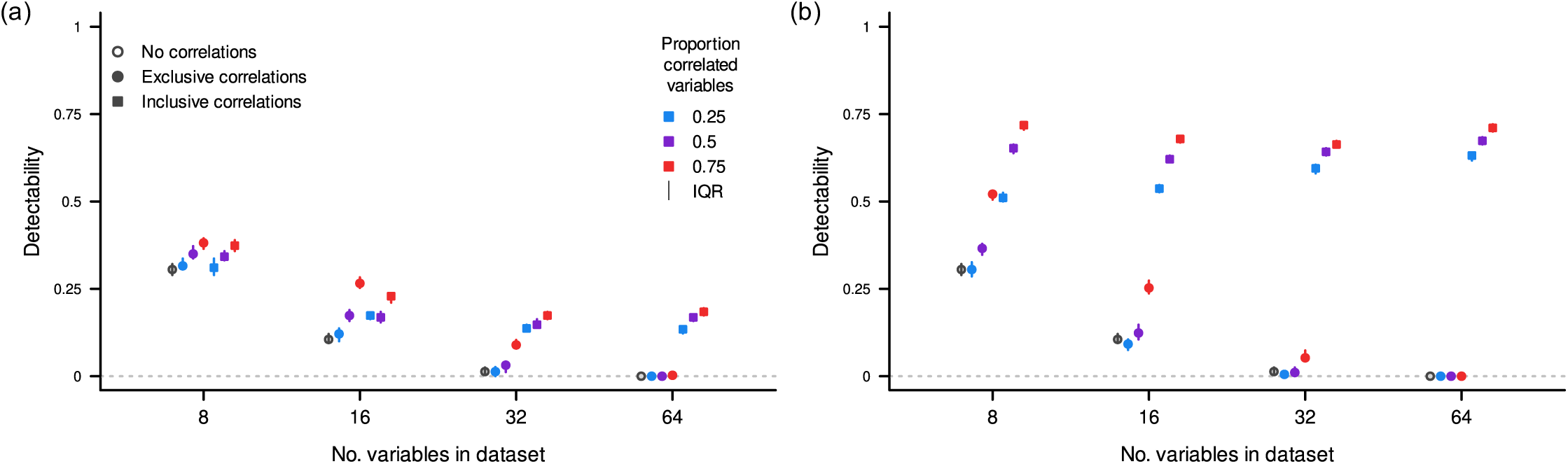
Influence of correlations between input variables (simulation set 2) on detectability of differences in ecosystem service (ES) values between archetypes. This example illustrates results from simulations where 20 archetypes were generated using K-means clustering, where input dataset variables shared (a) weak, and (b) strong correlations with each other. The ES values were simulated using a linear relationship with the predictor variable. Open circles represent detectability when none of the input variables were correlated. Filled circles represent simulations where the ES predictor variable was not correlated with other variables in the input dataset (exclusive correlations treatment). Filled squares represent simulations where the ES driver variable was correlated with other variables in the input dataset (inclusive correlations treatment). Each point is the computed median detectability from 50 replicate simulations, and vertical lines represent detectability inter-quartile ranges (IQR).

### 3.3. Set 3 – Influence of input variables with bounded distributions

Detectability increased as input datasets comprised increasing proportions of bounded variables (simulation set 3; Fig 3). However, the magnitude of this improvement declined as dataset size increased, such that improved detectability in datasets with a large number of variables was only apparent when over a quarter of the dataset comprised bounded variables. Our simulations also suggest that detectability was influenced by the interaction between the number and proportion of variables within a dataset that show bounded distributions (Supplementary Fig S3). Similar to results for simulations in sets 1 and 2, detectability improved when a greater number of archetypes were generated, declined for non-linear relationships between ES and input variables, and was not influenced by methodology used to develop archetypes.

**Fig 3.**
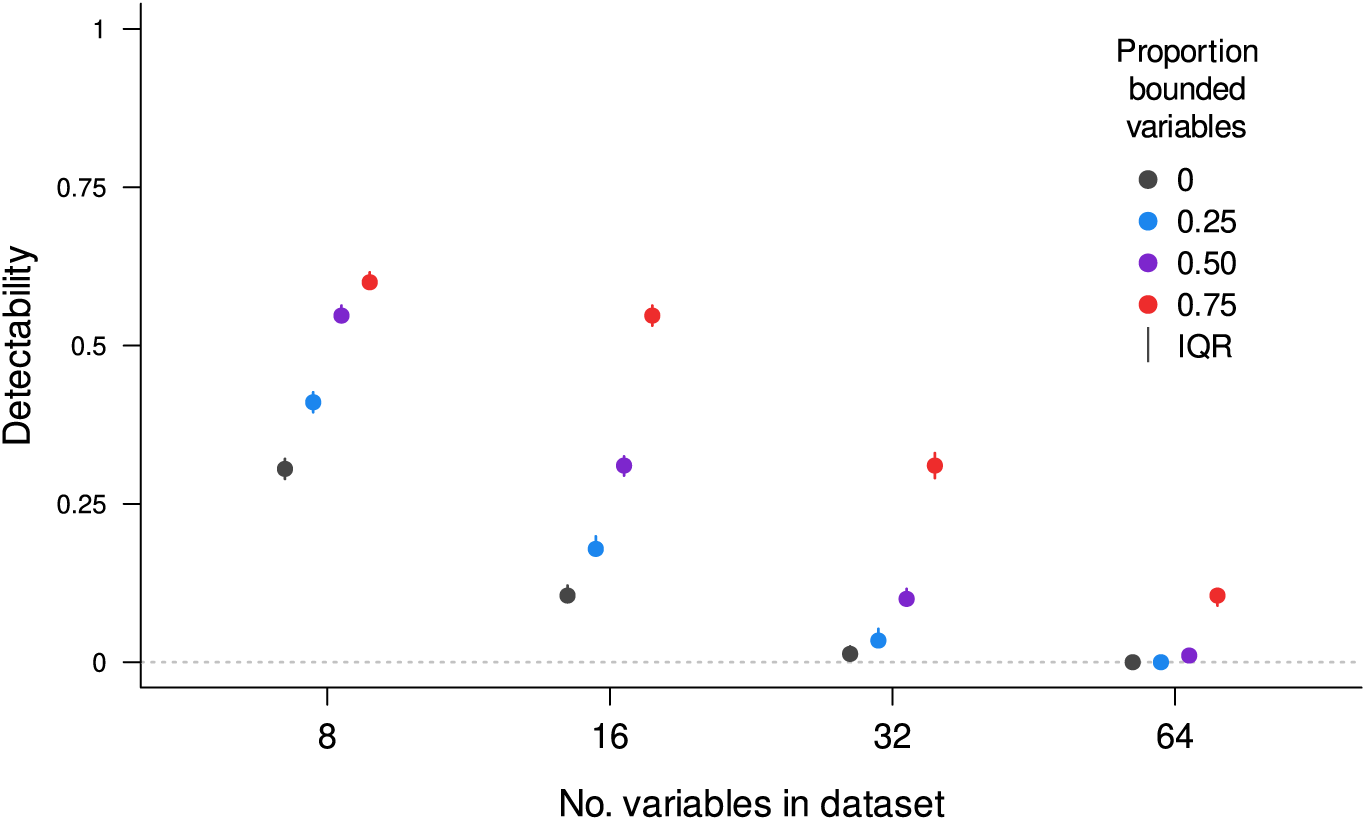
Effect of including input variables with bounded distributions (simulation set 3) on detectability of differences in ecosystem service (ES) values between archetypes. This example illustrates results from simulations where 20 archetypes were generated using K-means clustering. The ES values were simulated using a linear relationship with the predictor variable. Each point is the computed median detectability from 50 replicate simulations, and vertical lines represent detectability inter-quartile ranges (IQR).

### 3.4. Set 4 – Influence of input variables with multi-modal distributions

Inclusion of variables with multi-modal distributions (simulation set 4) had a strong influence on detectability, which displayed very high sensitivity to whether the ES predictor variable was multi-modal (inclusive treatment) or not (exclusive treatment). Small and large datasets containing even a few exclusive multi-modal variables showed large declines in, or no detectability (Fig 4). Datasets containing inclusive multi-modal variables showed substantial increases in detectability. However, detectability in these cases declined as the datasets included an increasing number of multi-modal variables. This decline in detectability was largely driven by the increasing number of multi-modal variables in datasets, though the interaction between the number and proportion of such variables does influence detectability when considering small datasets with very few multi-modal variables (Supplementary Fig. S4). Consistent with results from other simulation sets, detectability declined with increasing dataset sizes, improved where greater number of archetypes were generated, declined for non-linear relationships between ES and input variables, and was not influenced by methodology used to develop archetypes.

**Fig 4.**
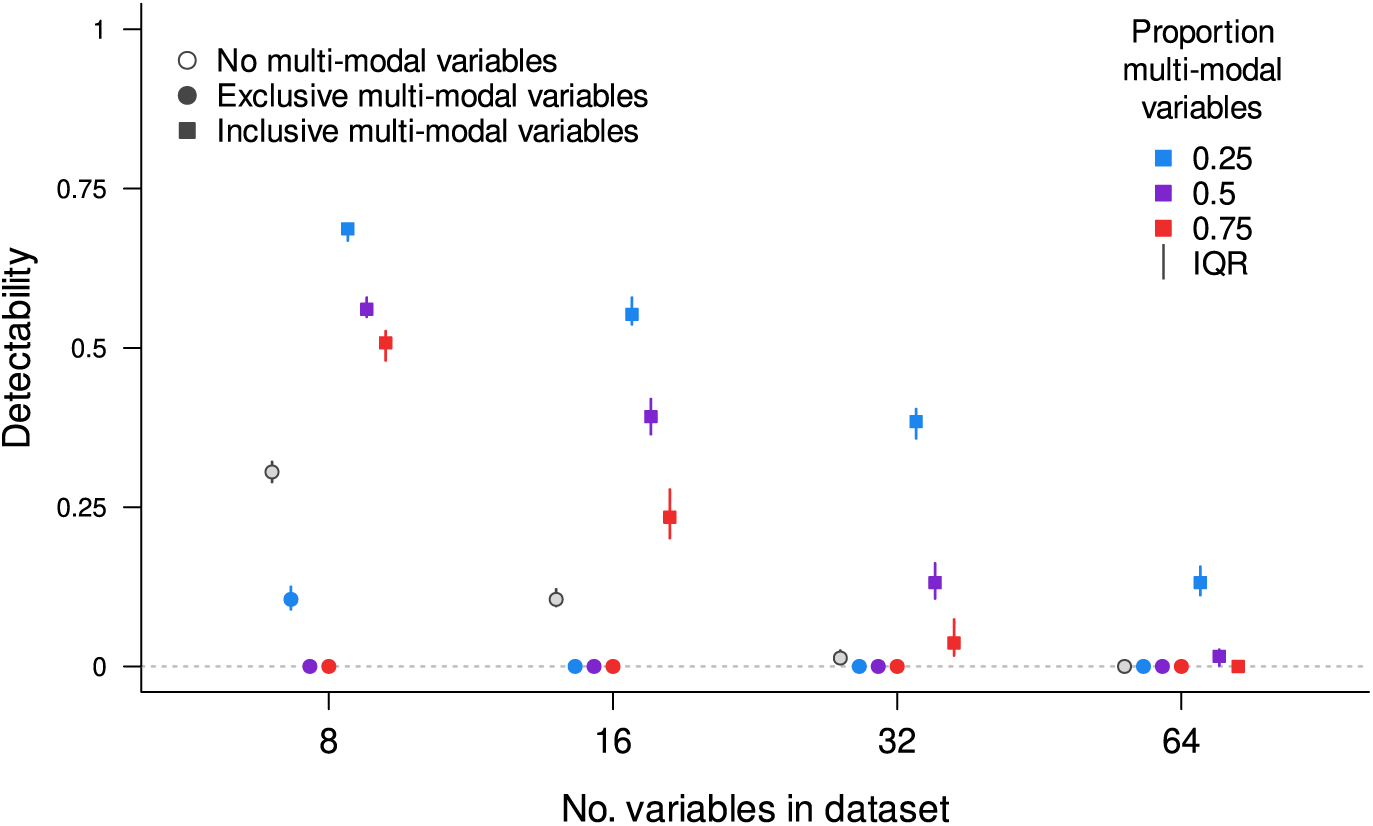
Effect of including input variables with multi-modal distributions (simulation set 4) on detectability of differences in ecosystem service (ES) values between archetypes. The results are illustrative of simulations where 20 archetypes were generated using K-means clustering. The ES values were simulated using a linear relationship with the predictor variable. Simulations where there were no multi-modal variables in the input dataset are represented by open circles. Filled circles represent simulations where multi-modal variables were included, but the ES predictor variable had a unimodal distribution. Filled squares indicate simulations where the ES predictor variable also showed a multi-modal distribution. Each point is the computed median detectability from 50 replicate simulations, and vertical lines represent detectability inter-quartile ranges (IQR).

### 3.5. Relating archetype classifier performance and ES-archetype detectability score

Compiling results across all simulations, we observed large variation around a general negative relationship between distance of observations to their assigned archetype centroid (a simple measure of how well the classifier performed in identifying archetypes) and detectability. For example, if we considered all simulations where archetypes were generated using K-means clustering and the simulated ES was a linear function (Fig 5) we observed very high, as well as very low detectability across a range of median distances to archetype centroids. Exceptions to this were when distances were either very low (showing relatively high detectability) or very large (no detectability), which might drive the overall negative relationship. However, the composition of input dataset variables (correlation structure and variable types) could substantially influence detectability (Supplementary Fig S5). Hence, we conclude that better archetype classifier performance may be a poor predictor of detectability.

**Fig 5.**
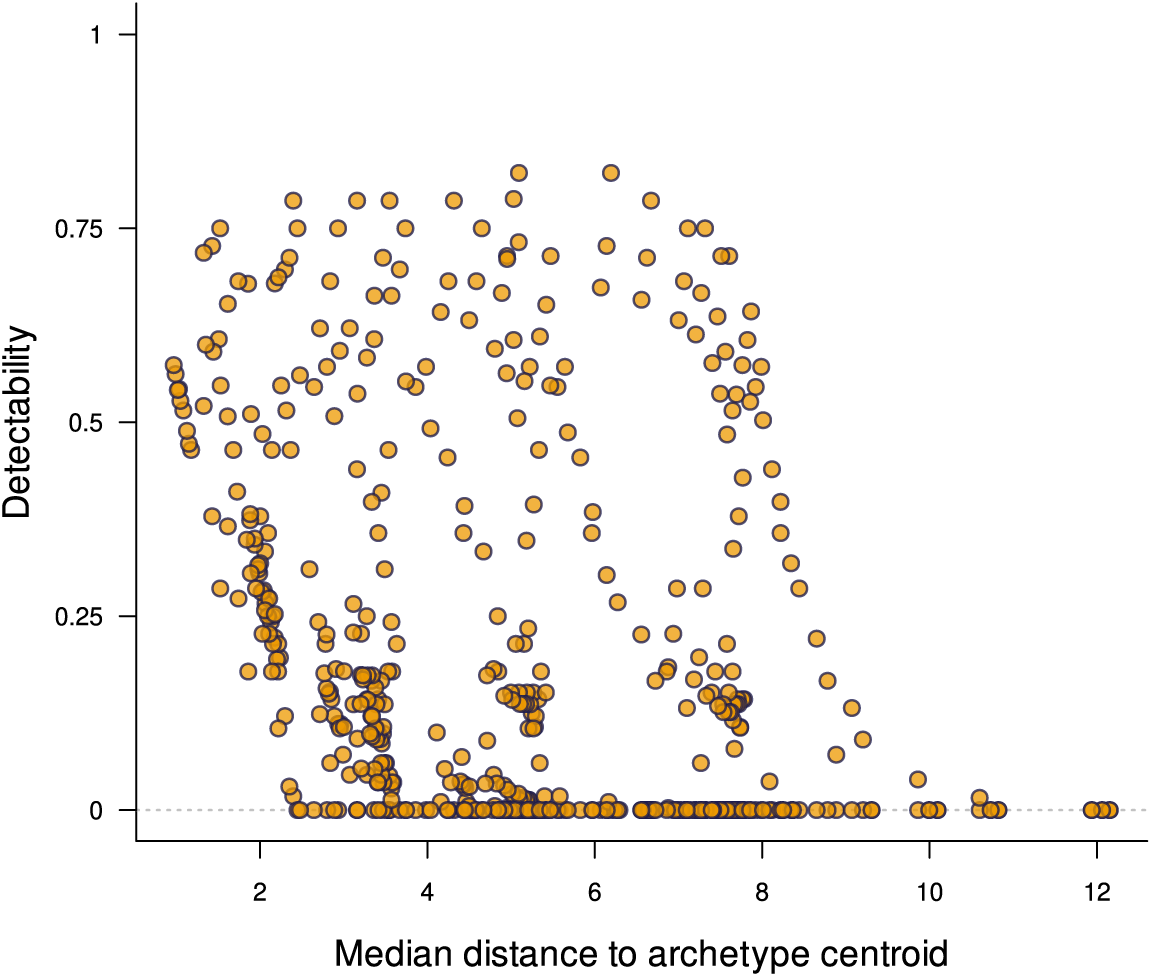
Change in detectability with increasing distance of observations to archetype centroids. Points represent median distances (50 replicates) from all simulations where ES values were simulated as a linear function of a predictor variable, and archetypes were generated using K-means clustering. See Supplementary Fig S5 for a detailed interpretation and decomposition by treatments.

## 4. Discussion

One of the clearest and most consistent observations in our simulations was that detectability of differences in ES values between archetypes declined with an increase in the number of variables in the input dataset. This result could represent an extension of what has been termed as the ‘curse of dimensionality’ (Bellman, 1957, 1961; Chandrasekaran and Jain, 1974; Trunk, 1979; Topchy and Punch, 2003; Gorban and Tyukin, 2018; Zollanvari, James and Sameni, 2020) – a commonly encountered challenge in machine learning and classification tasks. It describes the phenomenon where classification performance tends to decline as the number of variables within the dataset being classified becomes large and is a consequence of a fixed number of observations being distributed within an increasingly multi-dimensional volume determined by the number of input variables. In our simulations this manifests as an increase in the distance of observations to archetype centroids as the number of variables in datasets increase. In practical terms, larger distances between observations and archetype centres imply that, as the number of input variables increase, the archetype centroid becomes increasingly unrepresentative of the observations assigned to an archetype. Additionally, the contribution of the ES predictor variable to demarcating archetypes gets diluted by the large number of unrelated variables, resulting in lower detectability. One potential approach for mitigating the curse of dimensionality would require a disproportionate increase in the number of observations in a dataset to accommodate the increases in number of dataset variables. In socio-ecological archetype analyses this translates to the collation of datasets at finer spatial resolutions or increasing the extent of the study area to include more observations. Neither of these are feasible solutions given the general constraints of data availability for such analyses, and the fixed spatial extents over which analyses are required. Hence, while archetype analysis excels at distilling the complexity of data with many variables, post-hoc association analyses would see improved performance with a relatively restricted set of carefully chosen variables.

The correlation structure of the dataset, i.e., the relationships between input variables, can play an important role in mitigating the loss of detectability we observe with increasing dataset sizes. In general, our simulations show that an improvement in detectability can be expected when the ES predictor variable is correlated with other variables in the input dataset (inclusive correlation treatment). Detectability further improves as the proportion of correlated variables and strength of correlations increase. Even in simulations where the ES predictor variable was not amongst the correlated subset of variables (exclusive correlations), detectability could improve, but only if the input dataset comprised fewer variables. The case for fewer variables is also reinforced if we consider datasets where input variables were weakly correlated. Here, benefits of correlations between variables were influential when fewer correlated variables made up larger proportions of the input dataset. Filtering out correlated variables in datasets (e.g., by retaining only one from a pair of correlated variables) is a common pre-processing step in archetype analyses (Václavík *et al*., 2016; Rocha *et al*., 2020; Beckmann *et al*., 2022; Goodwin *et al*., 2022; Obringer and White, 2023), justified by the need to reduce collinearity in the data. Variables are screened using correlation coefficient thresholds, which differ between studies (Feng *et al*., 2019). However, archetype analyses strive to identify patterns of association in the data, rather than to ascribe causality or parametrise causal relationships where collinearity in the data can be an obstacle to interpretation (Dormann *et al*., 2013; Feng *et al*., 2019). Hence, post-hoc association analyses of archetypes may not be as sensitive to issues of collinearity, and our simulation results suggest liberal criteria (i.e., high correlation coefficient thresholds) could be employed to improve performance. The decision process to include/exclude correlated variables, and the thresholds to use can seem subjective, requiring a pragmatic approach. For example, in the identification of European agri-environmental archetypes, Beckmann *et al*. (2022) justified retaining highly correlated soil compositional variables, as well as elevation and terrain indices due to their established influence on ecosystem processes. Such an approach represents the continued use of expert domain knowledge throughout the process of developing archetypes, and is accepted as an important strategy to improve classifier performance and downstream application (Wardropper *et al*., 2016; Oberlack *et al*., 2019; Wicki *et al*., 2023; Pratzer *et al*., 2024).

Evaluating the influence of variables with non-unimodal distributions on classification of socio-ecological data and further downstream analyses is relatively unexplored. This is unexpected given the ubiquity of such variables (also binary, nominal and ordinal variables) in ecological datasets. Variables exhibiting bounded distributions—a special case of bimodal distributions truncated within a range—include any that can be expressed as a percentage or proportion. Hence, they encompass a diverse set of socio-ecological variables, such as, percentage composition of soil, the proportion of distinct land-cover types aggregated at specific (coarser) spatial resolutions, demographic composition, the contribution of specific economic activities to income generation, etc. Variables displaying multi-modal distributions, represented in our simulations as variables with three distinct modes, could emerge in ecological datasets as consequences of sharp geographic discontinuities or boundaries. An illustrative example is the occurrence of abrupt changes in bedrock material over short distances, with knock-on effects on other ecologically relevant variables (Hahm *et al*., 2014; Augusto *et al*., 2017; Callahan *et al*., 2022). Our simulations demonstrated that the inclusion of bounded variables can provide a net increase in detectability as the proportion of bounded variables within a dataset increase, especially if the total number of input variables is restricted. However, incorporating variables with multi-modal distributions in post-hoc association analyses can lead to erratic outcomes. Detectability of differences in ES-archetype associations improves only when the ES predictor variable also exhibits a multi-modal distribution (i.e., inclusive treatment), and when the total number of multi-modal variables in the dataset is low. Detectability substantially declines if the ES predictor is not itself a multi-modal variable (i.e., exclusive treatment). It is important to note that post-hoc association analyses assume that archetypes are generated with little-to-no prior knowledge of the ESs that will be used downstream. Consequently, the inclusion of multi-modal variables in the input dataset poses a risk to the usefulness of the archetypes that will be subsequently generated.

As anticipated, we observed reduced detectability when ES-predictor relationships showed strong non-linearity compared to linear and weakly non-linear relationships. Detectability between linear and weakly non-linear relationships did not differ. Conversely, we noted that detectability increased as we increased the number of archetypes generated from input datasets. This is an expected outcome as generating more archetypes partitions the data space into a greater number of classes, thereby, reducing within-class variability. While finding an objective answer to if, and how many clusters exist within a dataset remains a challenge (Kaufman and Rousseeuw, 2009; Eisenack *et al*., 2019; Sietz *et al*., 2019; Rocha *et al*., 2020; Wicki *et al*., 2023), it is crucial to stress that increasing the number of generated archetypes to enhance performance of post-hoc associations analyses would be counterproductive. Meaningful improvements in detectability would require the generation of a very large number of additional archetypes, compromising a key strength of archetype analyses – the reduction of dataset complexity, which is important for decision support and effectively communicating decision making frameworks to stakeholders. In our simulations we generated a maximum of 20 archetypes, which falls within the reported range of studies reviewed by Oberlack *et al*. (2019).

We explored two methods for classifying input datasets into archetypes: K-means clustering and SOMs. Notably, these methods exhibited virtually identical performance, and detectability did not differ between the various grid configurations within SOMs for a given number of required archetypes. Furthermore, while a general negative relationship between detectability and the distance of observations from archetype centroids exists, there is also substantial variability around this relationship. This underscores that for a given real-world dataset, the methodological efforts to identify effective archetypes do not guarantee superior performance in post-hoc association analyses. In other words, good archetypes can exhibit poor associations with ESs, while seemingly less effective archetypes can exhibit strong associations. Consequently, our findings suggest that efforts to fine-tune and choose between these two common archetype classification methods may not be the most productive approach. Instead, a greater focus should be on establishing criteria for identifying a limited (suitable) set of input variables with specific desirable characteristics (e.g., correlation structure, types of distributions), from which archetypes can be derived to provide greater utility in subsequent analyses. This approach would involve three steps. First, a thorough consultation with stakeholders and potential end-users to identify ESs of interest, and crucially, to define acceptable minimum levels of detectability. Second, the collation of data for a comprehensive set of potential ES predictor variables that incorporates expertise of domain specialists. Lastly, selecting a subset of these variables as inputs to generate archetypes. In this final step simulated ES variables could be used in conjunction with observed ES variables, if available. Variable selection could follow the procedures presented in this study or broadly adhere to principles of feature extraction (Hilario and Kalousis, 2008; Reddy *et al*., 2020; Tsai, Baldwin and Gopaluni, 2021), that can improve the performance of machine learning models.

## 5. Conclusion

The formulation of socio-ecological archetypes intended to have a general utility for decision making against multiple sustainability indicators now, and in the future, would be a powerful tool. However, this approach assumes that sustainability metrics can be partitioned by archetype identity into distinct value distributions. Our simulations identify constraints to this assumption, highlight key challenges in linking archetypes to sustainability metrics in post-hoc analyses, and suggest options for mitigation. We argue that rather than focussing on classifier performance, methodological choices should equally prioritise how the generated archetypes may relate to sustainability indicators in subsequent analyses, even when the indicators are unknown at the outset. The usability of the archetype framework for decision support could be enhanced by (1) limiting the number of input variables when classifying archetypes (reducing the curse of dimensionality); (2) using liberal correlation coefficient thresholds when selecting input variables (due to low sensitivity to collinearity); (3) avoiding the generation of a large number of archetypes (only provides marginal improvement in analytical performance); and (4) identifying the presence and influence of bounded and multi-modal input variables on down-stream analyses (can lead to erratic outcomes). The resulting archetypes would also be more adaptable to shifts in response variables of interest. This, in turn, ensures the consistency of the generated archetypes, which is important for stakeholder confidence in a decision support tool, as well as for monitoring decision outcomes that unfold over extended time horizons. Future research should explore methods, such as fuzzy archetype classification (Rao and Srinivas, 2006; Cullum *et al*., 2017; Eisenack *et al*., 2019), to mitigate the limitations of discrete archetypes and leverage, rather than lose, information on within-archetype variation. Such an approach could offer a more flexible decision support framework, capable of tackling sustainability challenges at both coarse and fine scales.

## Code availability

Code to run simulations and analyses are available on request from the corresponding author (VV).

