## Supplementary Fig for "Sensitivities in associating land-system archetypes with sustainability metrics: Insights from simulations"

---

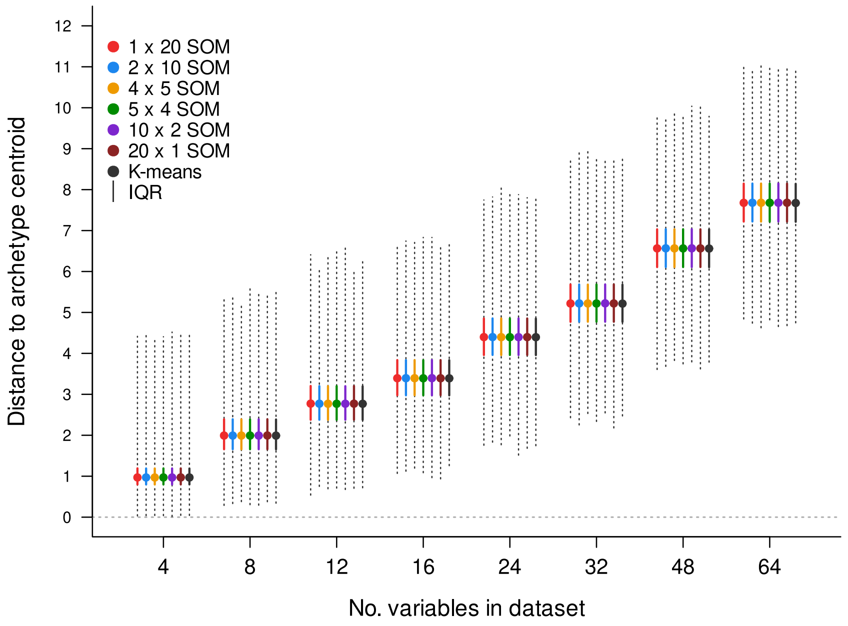


**Fig S1.** Increase in distance of data points to archetype centres with an increase in the number of variables in input datasets used to generate archetypes. In this specific example, 20 archetypes are generated using SOMs (and all possible grid topologies) and K-means clustering. Points represent median distance values using 50 replicate simulations. Vertical solid lines represent distance interquartile ranges, and dotted vertical lines represent the range of distance values observed in simulations. This illustrates the identical behaviour of K-means clustering and SOMs and the different SOM grid topologies. Qualitatively, this pattern of increasing distance with an increase in number of input variables is obtained regardless of the number of archetypes being generated.

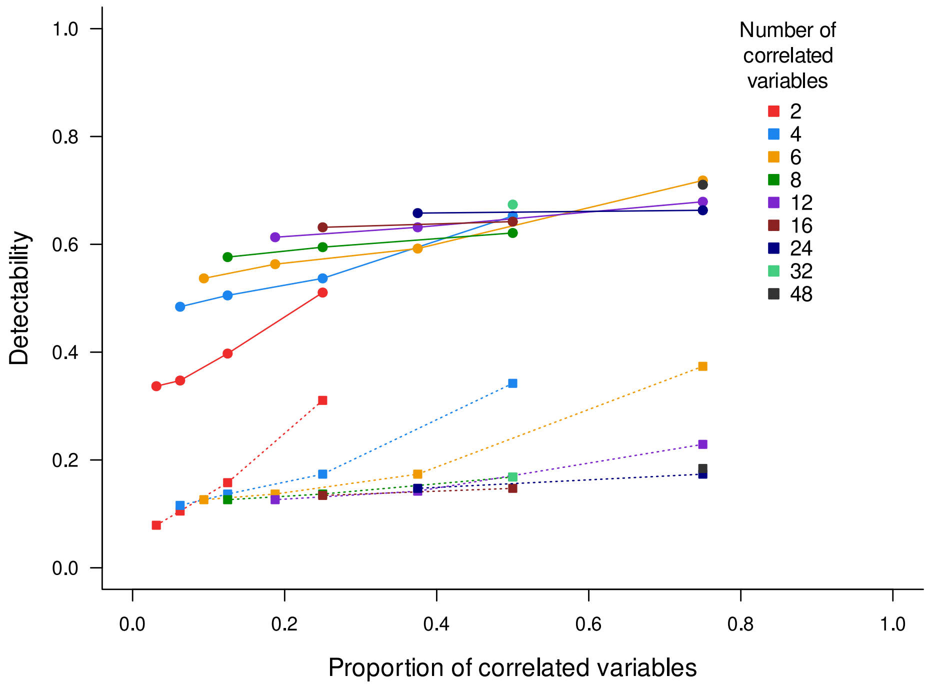


**Fig S2.** The interaction between number and proportion of correlated variables within input datasets on detectability of differences in simulated ES values between archetypes. This example is for an ES that is a linear function of the predictor variable, and where 20 archetypes are being generated from the input dataset using K-means clustering. Filled circles with solid lines represent simulations where strong correlations exist between input variables. Filled squares connect by dotted lines represent simulations where weak correlations exist between input variables.

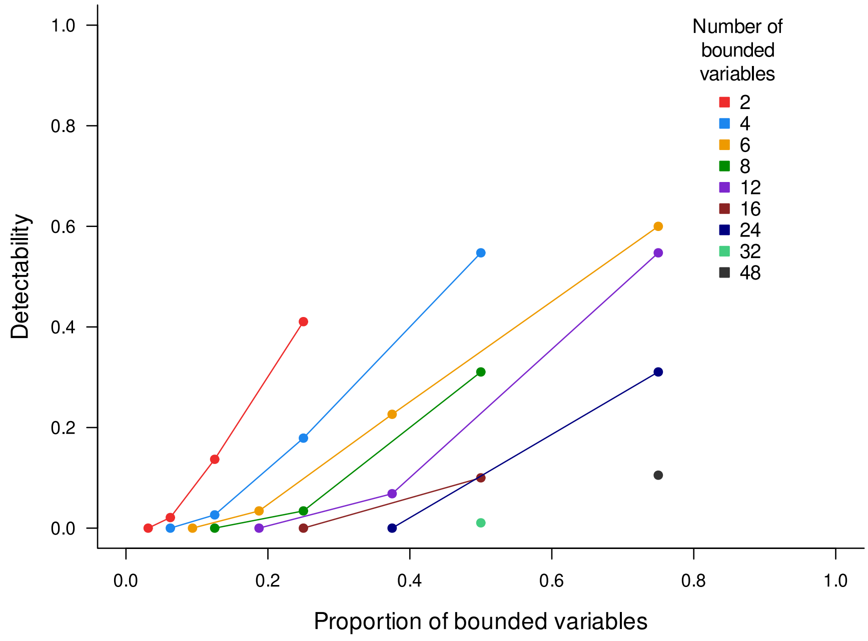


**Fig S3.** The interaction between number and proportion of bounded variables within input datasets on detectability of differences in simulated ES values between archetypes. This example is for an ES that is a linear function of the predictor variable, and where 20 archetypes are being generated from the input dataset using K-means clustering.

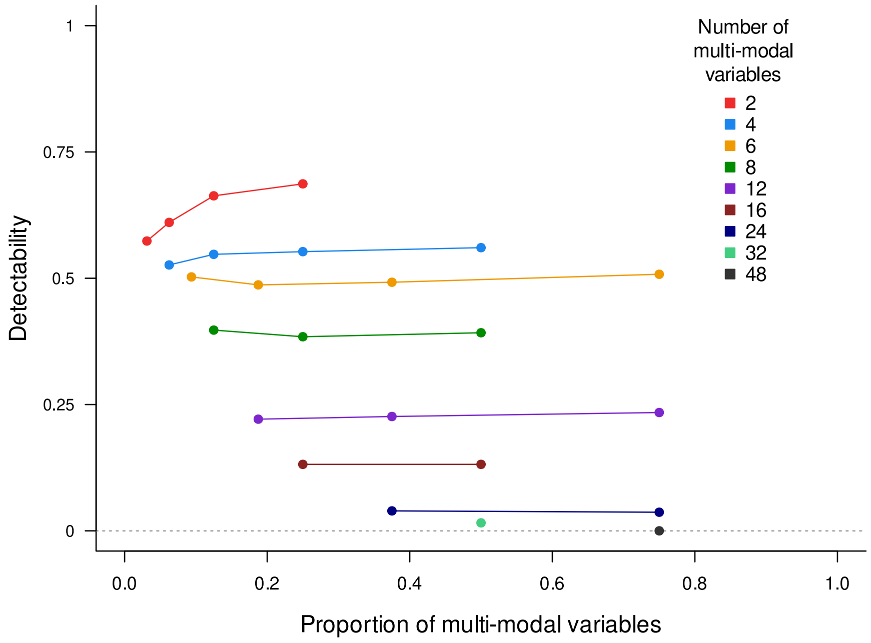


**Fig S4.** The interaction between number and proportion of multi-modal variables within input datasets on detectability of differences in simulated ES values between archetypes. This example is for an ES that is a linear function of the predictor variable that also exhibits a multi-modal distribution (inclusive multi-modal treatment), and where 20 archetypes are being generated from the input dataset using K-means clustering.

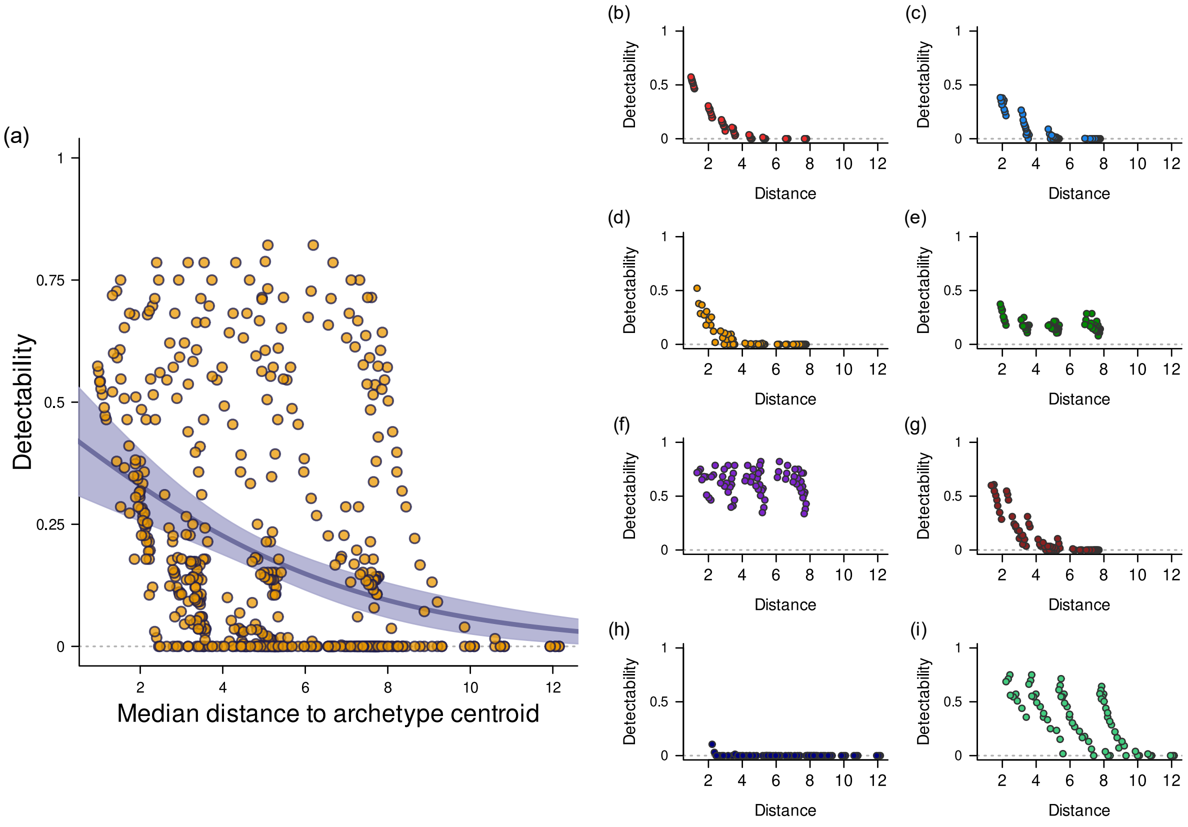


**Fig S5.** Relationship between detectability and distance of data points from archetype centres. Data presented here are for simulation scenarios where the simulated ES is a linear function of the predictor variable; archetypes are generated using K-means clustering; and simulation outputs for all levels of the 'number of archetypes' treatment are combined. Each point in the figures represent the median detectability value (from 50 replicates) of individual simulation treatment combinations. The complete dataset is presented in (a), where a generalised linear model with binomial errors identifies a negative relationship between detectability and distance (solid regression line with shaded 95% confidence intervals). Panels (b) to (i) visualise individual treatments within (a), where input datasets contain (b) uncorrelated and normally distributed variables; (c) – exclusive and weakly correlated variables; (d) – exclusive and strongly correlated variables; (e) – inclusive and weakly correlated variables; (f) – inclusive and strongly correlated variables; (g) – bounded variables; (h) exclusive, multi-modal variables; and (i) – inclusive, multi-modal variables. Results from individual treatments, especially (e), (f), (g) and (i) suggest that the composition of the input dataset (e.g., the relationship between variables and variable type) can override the general negative relationship between detectability and distance in (a). Therefore, methodological choices to generate archetypes that only prioritise reduction in distance between observations and archetype centres may not lead to desired outcomes in post-hoc associations analyses (i.e., increased detectability).
