## supplementary file SF1 for "Sensitivities in associating land-system archetypes with sustainability metrics: Insights from simulations": readme.txt

Column description for file SF1_archetype_simualtion_results_table.csv

author: Varun Varma

Notes: Supplementary datafile SF1 for preprint - Varma et al. - "Sensitivities in associating land-system archetypes with sustainability metrics: Insights from simulations". See preprint for methods that generated these data.

version date: 01-08-2024
------------------------------------------------------------------


sno: sequence number - for identification.

sim_set: simulation set

description: main parameters being modified in the simulation set

es_type: functional form of ecosystem service relationship to predictor variable

cluster_type: method used to derive archetypes from input dataset. Either SOM or K-means

no_arch: Number of archetypes generated from the input dataset

n_row: if clustering type is SOM, this column specifies number of rows in the SOM grid topology  (NA if clustering type is K-means).

n_col: of clustering type is SOM, this column specifies the number of rows int he SOM grid topology (NA if clustering type is K-means).

n_vars: number of input variables in the dataset used to generate archetypes

n_var_cor: number of correlated variables in the input dataset.

prop_var_cor: proportion of bounded variables in the input dataset.

cor_level: If there are correlated variables in the input dataset, this column specifies the level of correlation (correlation coefficient) between variables

n_var_bound: number of variables with bounded distributions in the input dataset.

prop_var_bound: proportion of variables with bounded distributions in the input dataset.

n_var_mmodal: number of variables with multi-modal distributions in the input dataset.

prop_var_mmodal: proportion of variables with multi-modal distributions in the input dataset.

inc_exc: valid only for set 2 (correlated input variables) and set 4(multi-modal input varaibles) simulations. Indicates if the treatment is inclusive (inc) or exclusive (exc) of the ecosystem service driver variable. For example - for set 2 simulations, 'inc' indicates if the ecosystem service driver variables was also correlated to other variables in the input dataset.

det_med: median detectabiltiy from 50 replicates

det_loq: lower quartile of detectability (50 replicates)

det_upq: upper quarlite of detectability (50 replicates)

dist_med: median distance of observations to archetypes centers (50 replicates)

dist_loq: lower quartile of distance of observations to archetype centers (50 replicates)

dist_upq: upper quartile of distance of observations to archetype centers (50 replicates)

dist_min: minimum distance of observations to archetype centers

dist_max: maximum distance of observations to archetype centers
